# Hyperoxia improves repeated-sprint ability and the associated training load in athletes

**DOI:** 10.1101/2021.11.22.469617

**Authors:** Shannon Cyr-Kirk, François Billaut

## Abstract

This study investigated the impact of hyperoxic gas breathing (HYP) on repeated-sprint ability (RSA) and on the associated training load (TL). Thirteen team- and racquet-sport athletes performed 6-s all-out sprints with 24-s recovery until exhaustion (power decrement ≥ 15% for two consecutive sprints) under normoxic (NOR: F_I_O_2_ 0.21) and hyperoxic (HYP: F_i_O_2_ 0.40) conditions in a randomized, single-blind and crossover design. The following variables were recorded throughout the tests: mechanical indices, arterial O_2_ saturation (S_p_O_2_), oxygenation of the vastus lateralis muscle with near-infrared spectroscopy, and electromyographic activity of the vastus lateralis, rectus femoris and gastrocnemius lateralis muscles. Session TL (work x rate of perceived exertion) and neuromuscular efficiency (work/EMG) were calculated. Compared with NOR, HYP increased S_p_O_2_ (2.7 ± 0.8%, Cohen’s effect size ES 0.55), the number of sprints (14.5 ± 8.6%, ES 0.28), the total mechanical work (13.6 ± 6.8%, ES 0.30) and the session TL (19.4 ± 7.0%, ES 0.33). Concomitantly, HYP increased the amplitude of muscle oxygenation changes during sprints (25.2 ± 11.7%, ES 0.36) and recovery periods (26.1 ± 11.4%, ES 0.37), as well as muscle recruitment (9.9 ± 12.1%, ES 0.74) and neuromuscular efficiency (6.9 ± 9.0%, ES 0.24). We conclude that breathing a hyperoxic mixture enriched to 40% O_2_ improves the total work performed and the associated training load during an open-loop RSA session in trained athletes. This ergogenic impact may be mediated by metabolic and neuromuscular alterations.

## 2 Introduction

The nature and the magnitude of a training effect are dictated by the frequency, duration and intensity of the exercise, the so-called training load (TL). Those parameters are modulated multiple times during a training cycle to manage fatigue accumulation and ensure progressive, specific physiological adaptations. Enhancement in athletic performance is well known to be attributable to the controlled fluctuation of the TL throughout the year (Busso, Denis, Bonnefoy, Geyssant, & Lacour, 1997; Fitz-Clarke, Morton, & Banister, 1991). Therefore, the understanding and quantification of the impact of varied stimuli on TL represent a key element to sport performance optimization (Foster et al., 2001).

Among the multiple stimuli used by athletes and coaches to increase TL and promote physiological adaptations, oxygen supplementation (i.e., hyperoxia) appears very attractive due to nearly all physiological functions relying on this gas. With growing popularity since its approbation by the World Anti-Doping Agency (WADA) in 2001, hyperoxia can be obtained by increasing the inspired oxygen fraction (F_I_O_2_, normobaric hyperoxia) and/or the barometric pressure (hyperbaric hyperoxia). Performance during aerobic exercise such as time trials, time to exhaustion, graded exercise tests and dynamic muscle function tests is, not surprisingly, acutely improved under hyperoxic conditions(Adams & Welch, 1980; Amann et al., 2006; Peltonen et al., 1997; Richardson et al., 1999). Those improvements are mainly derived from the increase in arterial hemoglobin saturation (S_a_O_2_), arterial content of oxygen (C_a_O_2_) and, therefore, the systemic O_2_ delivery to organs and skeletal muscles (Peltonen et al., 1995; Powers, Martin, & Dodd, 1993), which attenuate exercise-induced hypoxemia (Nummela, Hämäläinen, & Rusko, 2002) and alter energy metabolism (Linossier, Dormois, Arsac, Denis, & Lacour, 2000). Hyperoxia has also been used during repeated high-intensity exercise and results are promising. Higher peak and mean power output have been documented when breathing 100% O_2_, compared to room air, during two 30-s cycle sprints separated by 4 min (Kay, Stannard, & Morton, 2008), during five sets of 40 high-intensity breast strokes (~50 s) (Sperlich et al., 2011), and during ten 15-s cycling sprints interspersed with 45 s of recovery (Porter, Reed, & Jones, 2020). Overall, these data indicate that hyperoxia may be beneficial to exercise performance in varied sport settings, although the activity patterns of intermittent sports such as rackets and team sports have received limited attention so far.

During most team and racket sports, athletes repeat brief maximal or near-maximal efforts (i.e., sprints, changes of direction, jumps) with short recoveries at low to moderate intensity, throughout the duration of the competition. This physical quality, called repeated-sprint ability (RSA), is critical to success, and coaches and support staff use different training methods to enhance RSA during training (Billaut & Bishop, 2009; Bishop, Girard, & Mendez-Villanueva, 2011; Girard, Mendez-Villanueva, & Bishop, 2011). Many metabolic (e.g. phosphocreatine resynthesis, aerobic and anaerobic glycolysis, metabolite accumulation) and neuromuscular factors (e.g. muscle activation strategies) determine RSA, which have been reviewed in details elsewhere (Billaut & Bishop, 2009; Bishop et al., 2011; Girard et al., 2011). While the short maximal efforts rely heavily on anaerobic metabolic systems to produce ATP as rapidly as possible during the first sprints, early research using muscle biopsy has also well demonstrated that when sprints are repeated with brief and incomplete recovery, the contribution of the anaerobic glycolysis to ATP resynthesis diminishes and the aerobic metabolism becomes more critical (Gaitanos, Williams, Boobis, & Brooks, 1993). This is confirmed by studies using hypoxia to reduce tissue oxygen availability. For example, Billaut et al. demonstrated with near-infrared spectroscopy (NIRS) that reducing arterial O_2_ saturation led to a decline in muscle oxygenation which impaired RSA in trained athletes (Billaut & Buchheit, 2013). Furthermore, the recovery of performance between sprints is purely fueled via aerobic energy sources, and muscle tissue reoxygenation has been shown to be another determinant of RSA (Billaut & Buchheit, 2013; Buchheit & Ufland, 2011). With such metabolic changes occurring during a repeated-sprint training session, one may expect a hyperoxic mixture to enhance some the aerobic components of RSA and, thereby, better maintain performance. A higher exercise intensity or the possibility to perform more sprint repetitions during a given training session would lead to a higher TL, and theoretically, greater training adaptations over time. However, to our best knowledge, no study has assessed the influence of HYP on RSA and the associated TL.

Before promoting such practice in training, one may first assess the efficacy of adding hyperoxia to a repeated-sprint training session with athletes. Therefore, we examined the effects of acute normobaric hyperoxia on RSA, key psycho-physiological parameters and TL in trained athletes. We hypothesized that hyperoxia would enhance muscle reoxygenation and better maintain muscle recruitment, which would lead to enhanced RSA and TL.

## 3 Materials and methods

### 3.1 Ethics approval

The study was approved by the Ethics Committee of University Laval, and adhered to the principles established in the Declaration of Helsinki. Participants provided written informed consent after being informed of experimental procedures, associated risks and potential benefits.

### 3.2 Participants

Thirteen athletes (including 5 women) volunteered for this study (mean ± SE; age, 23.0 ± 2.7 yr; body mass, 70.7 ± 14.9 kg; stature, 175.5 ± 7.3 cm; percent body fat, 14.00 ± 4.38 %). All participants were non-smokers, free of health problems, and did not use any medication or any other tobacco/nicotine products. Participants were training 8 (range 6-12) hour per week (in a sport that included high-intensity short intervals (e.g., basketball, hockey and badminton). They avoided vigorous exercise, drug and nutritional supplements 48 h before test and refrained from alcohol and caffeine for 12 h. They were also asked to consume a regular meal at least 2 h before every session and to keep a constant training load during the course of the study.

### 3.3 Experimental design

Participants visited the laboratory for one preliminary visit and two experimental trials during which all testing procedures were performed by the same investigator. Sessions were separated by a minimum of 2 days to avoid fatigue and a maximum of 7 days.

During the first visit, stature, body mass and percentage body fat (Tanita TBF-310; Tanita Corp. of America Inc., Arlington Heights, IL) were recorded. Resting heart rate and blood pressure (inclusion criteria, <140/100 mm Hg) were taken before trial in a seated position. The handlebars and seat were adjusted vertically and horizontally to each participants preference, and feet were secured to the pedals using toe clips. These settings were also replicated for all subsequent trials. Participants completed a 10-min standardized, specific RSA warm-up on a computer-controlled electrically braked cycle ergometer (Velotron Elite; RacerMate, Seattle, WA). After 3 min of passive rest, they performed a force-velocity (FV) test to determine the optimal cycling resistance (see *Force-Velocity Test* procedures below). Athletes had 5 min of passive recovery and were then familiarized with the RSA test procedures. They executed a shortened version of the RSA test (see *Repeated-Sprint Ability Test* procedures below) which included six 6-s sprints while wearing the breathing mask only.

Following the preliminary visit, athletes were randomised in a single-blind, cross-over design and asked to perform a RSA test in normoxia (NOR, F_I_O_2_ 0.21) and in normobaric hyperoxia (HYP, F_I_O_2_ 0.40). Normoxic and hyperoxic gas mixtures were delivered by a continuous-flow O_2_ concentrator (New Life Intensity 10, AirSep Corporation, Buffalo, NY) and administered via a rig of 3 x 170-liter Douglas bags connected to a facemask (AltitudeTech Inc, Kingston, ON, Canada). After the standardized warm-up, participants were instrumented with the NIRS device, electromyography electrodes and pulse oximeter. For both experimental trials, the breathing apparatus was installed one minute before the beginning of the RSA test and was removed after the last sprint. A 5-minute cool-down was observed after every trial.

### 3.4 Testing procedures

#### 3.4.1 Standardized warm-up

A standardised warm-up was realized by the athletes before the force-velocity and the RSA tests. It included a total of 3 min of moderate intensity cycling, two 15 s accelerations with 15 s active recovery, 1 min 30 s of moderate intensity and then three 6 s sprints at 80%, 90% and 100% of their maximal effort with 24 s of recovery between each of them. The last three minutes of the warm-up were passive rest where they could stay on the bike without pedaling or stand-up.

#### 3.4.2 Force-Velocity Test

Athletes performed five maximal 6-s cycle sprints against increasing resistances and interspersed with 4 min of passive recovery. Sprints were initiated with 5-s of cycling at a standardized cadence (between 80-90 rpm) with the same resistance of the following sprint. Athletes were allowed to stand-up for the first two pedal strokes and then remained seated for the duration of the sprint and were strongly encouraged to reach maximal power output as quickly as possible (same instructions were given for the RSA test). The individual power–velocity relationships were obtained, and the optimal cycling resistance was defined as the gear for which maximal cycling power output was reached.

#### 3.4.3 Repeated-Sprint Ability Test

After the standardised warm-up, participants performed consecutive maximal 6-s cycle sprints, separated by 24 sec of rest (19 sec of passive rest and 5 sec of progressive re-acceleration of the flywheel) until exhaustion on an electronically braked cycle ergometer (Velotron Elite, RacerMate, Seattle, WA). An electronic tablet with a visual and audible signal was placed directly in front of the participants for precise count-down. Participants were asked to stop pedalling immediately at the end of the sprint. The resistance determined during the force-velocity test was set at the beginning of the RSA test and did not change afterwards. Following the test, all instrumentation was removed, and participants performed a self-paced cool down of at least 3 minutes.

To reduce the inter-subject variability in performance decrement and to enable a more standardized level of fatigue between the participants, both RSA tests were stopped when participants completed two consecutive sprints with a percent decrement score (S_DEC_) higher than 15% or if they reached 20 sprints (Morin, Dupuy, & Samozino, 2011). The S_DEC_ across the sprints was determined as the percent difference between total and ideal performance (Glaister, Howatson, Pattison, & McInnes, 2008), as follows: S_DEC_ = (100 x (total performance / ideal performance)) – 100, where total performance is the sum of the PPO from all sprints performed, and ideal performance is the highest PPO recorded multiplied by the number of sprints performed.

The Velotron ergometer provides instantaneous, average and peak values for power output, pedal frequency and speed at 23 Hz. The mechanical work (kJ) performed during every sprint was calculated by integrating the instantaneous power curve over the 6 sec of the sprint. The number of sprints, the peak power output (PPO) and the total mechanical work (W_tot_, the sum of the work from all sprints) were measured and calculated. We also calculated the sum of the mechanical work values for the common sprints (i.e., the sprints performed by every participant in the two conditions, W_com_). For example, if a participant performed 12 sprints in NOR and 14 sprints in HYP, only the first 12 sprints of each condition were taken for the W_com_ calculation.

### 3.5 Exercise physiological responses

#### 3.5.1 Rate of perceived exhaustion (RPE)

Perceived exhaustion (RPE_tot_), perceived limb discomfort (RPE_lim_) and perceived difficulty breathing (RPE_bre_) were recorded from a modified Borg CR10 scale after every sprint. Participants were asked to point the number associated with their perceived effort on a printed scale. The session rating of perceived exertion (sRPE) was also calculated and considered as the main variable to measure TL (Foster et al., 2001). Calculation: sRPE = (W_tot_ x average rates of perceived effort).

#### 3.5.2 Arterial O_2_ saturation (S_p_O_2_) and heart rate (HR)

S_p_O_2_ was estimated via pulse oximetry (Nellcor Bedside; Nellcor Inc. Hayward, CA) with adhesive optodes placed on the forehead. The reproducibility and validity (intraclass correlation coefficient = 0.99) for this method of measurement for saturations above 80% has been shown to be in good agreement with hemoglobin O_2_ when compared with arterial blood gas measurements (Romer et al., 2007). The S_p_O_2_ and the HR were recorded at rest, and immediately after every sprint and every recovery period. The lowest value of S_p_O_2_ recorded during the RSA test was considered as S_p_O_2min_.

#### 3.5.3 Near-infrared spectroscopy (NIRS)

A portable spatially-resolved, dual wavelength NIRS apparatus (PortaMon, Artinis Medical Systems BV, The Netherlands) was installed on the distal part of the right vastus lateralis muscle belly (approximately 15 cm above the proximal border of the patella), parallel to muscle fibers, to quantify changes in the absorption of near-infrared light by oxyhemoglobin/myoglobin (O_2_Hb/Mb) and deoxyhemoglobin/myoglobin (HHb/Mb). Skinfold thickness was measured at the site of the application of the NIRS (9.0 ± 4.6 mm) using a Harpenden skinfold caliper (British Indicators Ltd, West Sussex, Great Britain) during the familiarization session and was less than half the distance between the emitter and the detector (i.e., 20 mm). This thickness is adequate to let near-infrared light through muscle tissue (McCully & Hamaoka, 2000). The device was packed in transparent plastic wrap to protect it from sweat and fixed with tape. Black bandages were used to cover the device from interfering background light. The position of the apparatus was marked with an indelible pen for repositioning in the subsequent visit.

A modified form of the Beer–Lambert law, using two continuous wavelengths (760 and 850 nm) and a differential optical path length factor of 4.95, was used to calculate micromolar concentrations in tissue O_2_Hb/Mb, HHb/Mb and total hemoglobin/myoglobin ([THb/Mb] = [O_2_Hb/Mb] + [HHb/Mb]). The tissue saturation index (TSI = [O_2_Hb/Mb] / [THb/Mb]) was also calculated. Exercise data was normalised to a 1-min baseline recorded before the warm-up of visit 2 and 3 (defined as 0 μM). The NIRS data were acquired continuously at 10 Hz and then filtered using a tenth-order Butterworth low-pass filter with a 4 Hz cut-off frequency. Analysis of muscle oxygenation was limited to [HHb/Mb] because this variable is less sensitive than [O_2_Hb/Mb] to perfusion variations during contraction and recovery and is closely related to the venous O_2_ content (De Blasi, Cope, Elwell, Safoue, & Ferrari, 1993; Ferrari, Mottola, & Quaresima, 2004), and to TSI because this parameter is independent of near-infrared photon path length in tissue and reflects the dynamic balance between O_2_ supply and O_2_ consumption in the microcirculation (Van Beekvelt, Colier, Wevers, & Van Engelen, 2001). From the filtered signal, the maximal and minimal [HHb/Mb], [THb/Mb] and TSI were identified for every sprint/recovery cycle from a 5-sec moving average to identify the maximal upper and lower limits of the signals (Rodriguez, Townsend, Aughey, & Billaut, 2018). We further calculated the deoxygenation amplitude during sprints (Δ[HHb/Mb]_exercise_ and ΔTSI_exercise_) as the average difference between maximal and minimal [HHb/Mb] and TSI values for every sprint. Similarly, the reoxygenation amplitude during post-sprint recovery periods (Δ[HHb/Mb]_recovery_ and ΔTSI_recovery_) was calculated as the difference between maximal and minimal [HHb/Mb] and TSI values for every recovery period.

Normalized NIRS values were also represented as percentage of RSA test completion. As the fewer number of sprints realized by a participant was five, we divided data series in five segments to obtain beginning (0%), 25%, 50%, 75% and 100% of sprint completion, and calculated the averages of all parameters for every segment.

#### 3.5.4 Electromyography (EMG)

The EMG signals of the *vastus lateralis*, *rectus femoris* and *gastrocnemius lateralis* were recorded from the dominant lower limb via surface electrodes (Delsys, Trigno Wireless, Boston, MA) during every sprint. A careful preparation of the skin (shaving, light abrasion and cleaning with an alcohol swab) on the electrodes sites was made before every test. Electrodes were fixed longitudinally (aligned parallel to the underlying muscle fibre direction) over the muscle belly. The EMG signal was pre-amplified and filtered (bandwidth 12-500 Hz, gain = 1,000, sampling frequency 2kHz) and recorded with Delsys hardware (Bagnoli EMG System; Delsys, Inc., USA). During postprocessing, the onset and offset of activation of all EMG bursts in all muscles and every sprint were detected manually. The activity of each muscle was determined by calculating the integral of the EMG (iEMG) and the median power frequency (MPF) between the onset and the offset of the last 6 subsequent bursts of the sprint. Values of iEMG of every muscle were summed together and a new sum-iEMG parameter was used to represent the global muscle activity during sprints (Billaut et al., 2013; Billaut, Physiology, Nutrition, and, 2009; Smith & Billaut, 2010). The sum-iEMG for all the sprints (sum-iEMG_tot_) and for the common sprints (sum-iEMG_com_) were also calculated.

Neuromuscular efficiency (NME) was calculated for every sprint as the ratio of work to sum-iEMG (NME = work/sum-iEMG) and expressed for all the sprints (NME_tot_) and for the common sprints (NME_com_). A decrease in NME would indicate perturbations in muscle contractility (Bigland-Ritchie, 1981). Mechanical and EMG parameters are reported as raw data and as percent of the first sprint value. Normalisation has been used widely when the effects of fatigue on force production and EMG activity are studied (Bigland-Ritchie, 1981; Hunter, Critchlow, Shin, & Enoka, 2004).

### 3.6 Statistical Analysis

All data are reported as means ± SD, percentage of normalized values or percentage of change from NOR. The NOR and HYP differences were analysed using Cohen’s effect size (ES) ± 90% confidence limits and compared to the smallest worthwhile change (0.2 multiplied by the between-participant SD) (Batterham & Hopkins, 2005; Hopkins, Marshall, Batterham, & Hanin, 2009). Effect sizes were classified as small (>0.2), moderate (>0.5) and large (>0.8), and only likely effects (>75%) are reported (Batterham & Hopkins, 2005; Hopkins et al., 2009).

## 4 Results

### 4.1 Mechanical and perceptual measurements

Performance variables are presented in **Table 1** and **Figure 1**. The PPO reached was similar in both conditions, but HYP extended the number of sprints performed before exhaustion compared to NOR (difference between groups 14.5 ± 8.6%, ES 0.28). The total work performed over the entire series also increased in HYP (13.6 ± 6.8%, ES 0.30), but when the same number of sprints was considered in both NOR and HYP, the work in these common sprints did not change meaningfully (2.0 ± 1.7%, ES 0.05). Similar effects were observed when analyzing work to body mass ratio (J/kg) on W_tot_ (13.6 ± 6.8%, ES 0.27) and W_com_ (2.0 ± 1.7%, ES 0.04).

**Table 1.** Mean changes in performance and perceptual exercise responses in the repeated-sprint ability test in normoxia (NOR, FIO2: 0.21) or hyperoxia (HYP, FIO2: 0.40).

|  | NOR | HYP | Differences between groups |  |
| --- | --- | --- | --- | --- |
|  |  |  | %D | Cohen's ES |
| N<sub>sprints</sub> (AU) | 12.5 ± 5.6 | 13.8 ± 4.8 | 14.5 ± 8.6 | **0.28 ± 0.17** |
| PPO (W) | 428.2 ± 124.6 | 434.1 ± 121.2 | 1.6 ± 1.2 | 0.05 ± 0.04 |
| W<sub>tot</sub> (kJ) | 24.47 ± 9.10 | 27.65 ± 9.80 | 13.6 ± 6.8 | **0.30 ± 0.15** |
| W<sub>com</sub> (kJ) | 24.33 ± 9.20 | 24.91 ± 9.70 | 2.0 ± 1.7 | 0.05 ± 0.04 |
| RPE<sub>tot</sub> (AU) | 5.6 ± 0.9 | 6.1 ± 0.9 | 8.9 ± 5.9 | **0.47 ± 0.32** |
| RPE<sub>bre</sub> (AU) | 5.9 ± 1.0 | 6.3 ± 1.0 | 7.5 ± 7.3 | **0.40 ± 0.39** |
| RPE<sub>lim</sub> (AU) | 5.7 ± 0.8 | 6.0 ± 1.4 | 1.7 ± 9.9 | 0.09 ± 0.52 |
| RPE<sub>tot</sub> max (AU) | 7.9 ± 1.8 | 8.3 ± 1.70 | 6.5 ± 8.3 | 0.27 ± 0.34 |
| RPE<sub>bre</sub> max(AU) | 8.1 ± 1.7 | 8.6 ± 1.3 | 7.3 ± 6.3 | **0.31 ± 0.26** |
| RPE<sub>lim</sub> max(AU) | 8.3 ± 1.4 | 8.5 ± 1.5 | 2.3 ± 5.9 | 0.10 ± 0.25 |
| TL (AU) | 139.52 ± 60.49 | 169.60 ± 69.88 | 22.3 ± 9.6 | **0.38 ± 0.17** |

Abbreviations: %D; percentage difference between changes in NOR and HYP, ES; effect size, Probability %, percentage chances for HYP to be higher/similar/lower than NOR, N<sub>sprints</sub>; number of sprints performed, PPO; peak power output, W<sub>tot</sub>; total mechanical work of all sprints, W<sub>com</sub>; total mechanical work of common sprints, RPE<sub>tot</sub>; perceived exhaustion, RPE<sub>lim</sub>; perceived limb discomfort, RPE<sub>bre</sub>; perceived difficulty breathing, TL; training load
Data are presented as means ± SD. Cohen's effect size ± 90% confidence limits. Clear changes between conditions are indicated in bold.

**Figure 1.**
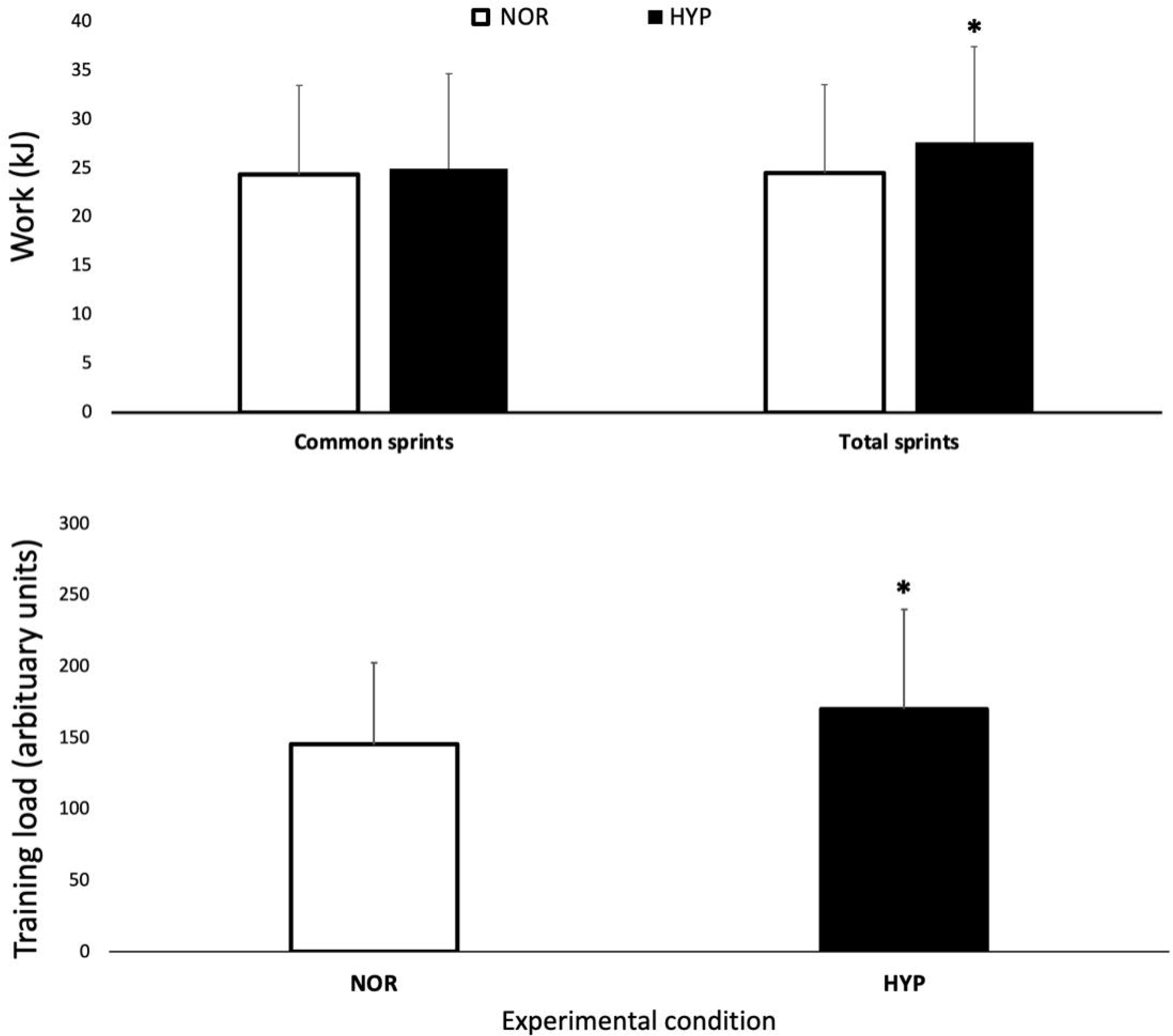
Mechanical work performed over the entire series of sprints (W_tot_) and for the same number of sprints performed in both conditions (W_com_) session training load over the repeated-sprint ability test in normoxia and hyperoxia (F_I_O_2_ 0.40). Data are presented as means ± SD. * Small effect between conditions. W_tot_ values were 24.47 ± 9.1 kJ in NOR and 27.65 ± 9.8 kJ in NYP. W_com_ values were 24.33 ± 9.2 kJ in NOR and 24.91 ± 9.7 kJ in HYP. TL was increased from 139.52 ± 60.49 AU in NOR to 169.60 ± 69.88 in HYP.

Importantly, HYP also increased the session TL by 19.4 ± 7.0% (ES 0.33). HYP also increased RPE_tot_ (8.9 ± 5.9%, ES 0.47) and RPE_bre_ (7.5 ± 7.3%, ES 0.40). There was no clear difference between conditions for RPE_lim_.

### 4.2 Arterial oxygenation and heart rate

Breathing the hyperoxic mixture increased the averaged S_p_O_2_ recorded during the entire series of sprints (2.7 ± 0.8, ES 0.55) as well as the lowest S_p_O_2_ (HYP 99.2 ± 0.8 vs NOR 95.1 ± 2.1, difference 4.4 ± 1.0, ES 1.85). Furthermore, HR was lower in HYP (HYP 163.1 ± 14.2 bpm vs NOR 164.8 ± 12.6 bpm, difference −1.1 ± 1.9, ES −0.24).

### 4.3 Muscle oxygenation

**Figure 2** displays the changes in NIRS variables over the RSA test in both NOR and HYP. During the sprints, the maximal metabolic alterations ([HHb/Mb]_max_ and TSI_min_) were not different between conditions, however we observed greater average muscle deoxygenation amplitude when sprints were performed in HYP compared with NOR (Δ[HHb/Mb]_exercise_: 25.2 ± 11.7%, ES 0.36 and ΔTSI_exercise_: 14.0 ± 13.3%, ES 0.62).

**Figure 2.**
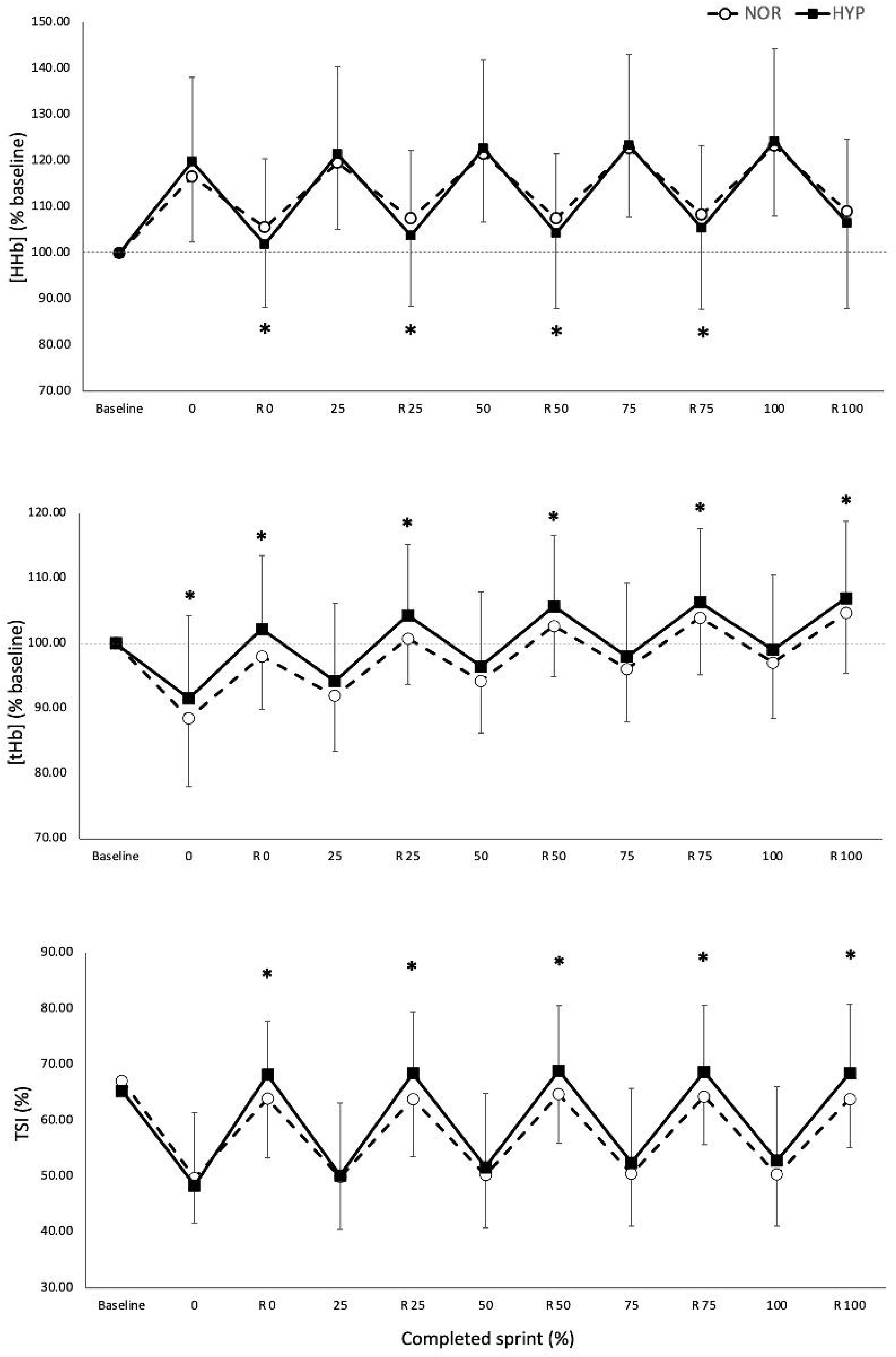
Maximal and minimal values of normalized deoxyhemoglobin/myoglobin concentration ([HHb/Mb], panel A), total hemoglobin/myoglobin concentration ([tHb/Mb], panel B) and tissue saturation index ([TSI], panel C) over the sprints and recovery periods for the five percentages of test completion in normoxia and hyperoxia (F_i_O_2_ 0.40). Data are presented as means ± SD, expressed as a percent of the baseline. * Small effect between conditions.

The metabolic status of the recovery periods between sprints was also altered by the condition. Muscle reoxygenation was greater in HYP (Δ[HHb/Mb]_recovery_: 26.1 ± 11.4%, ES 0.37 and ΔTSI_recovery_: 27.8 ± 15.7%, ES 0.53). This led to lower values of [HHb/Mb]_min_ in HYP at the beginning (−3.5 ± 4.1%, ES –0.25), 25% (−3.5 ± 5.2%, ES –0.24), 50% (−3.2 ± 5.1%, ES –0.22) and 75% (−3.0 ± 5.5%, ES –0.21) of test completion. Furthermore, TSI_max_ was higher in HYP at the beginning (7.0 ± 6.7%, ES 0.36), 25% (7.0 ± 6.9%, ES 0.36), 50% (5.8 ± 7.8%, ES 0.30), 75% (5.9 ± 7.8%, ES 0.31) and 100% (6.3 ± 7.9%, ES 0.33) of test completion. An increase in [THb/Mb]_max_ was also observed at the beginning (4.3 ± 4.7%, ES 0.45), 25% (3.3 ± 3.9%, ES 0.35), 50% (2.7 ± 3.5%, ES 0.29), 75% (2.1 ± 3.4%, ES 0.22) and 100% (1.9 ± 3.5%, ES 0.21).

### 4.4 Electromyographic activity

HYP increased sum-iEMG_tot_ by 9.9 ± 12.1% (ES 0.74) and sum-iEMG_com_ by 10.2 ± 12.0% (ES 0.77). When looking at muscles individually, HYP increased the average value of the *vastus lateralis* sum-iEMG_tot_ by 10.3% ± 11.8% (ES 0.71) and that of the *rectus femoris* sum-iEMG_tot_ by 14.8% ± 16.7% (ES 0.93), while no change was observed in the *gastrocnemius lateralis*.

NME values are presented in **Figure 3**. Mean value of NME_tot_ increased in HYP (6.9 ± 9.0%, ES 0.24) as well as mean value of NME_com_ (7.8 ± 8.7%, ES 0.27). Finally, MPF decreased from 82.3 ± 13.9 Hz to 77.85 ± 10.81 Hz in NOR and from 81.75 ± 13.22 to 78.14 ± 10.61 Hz in HYP, without showing any clear difference between conditions.

**Figure 3.**
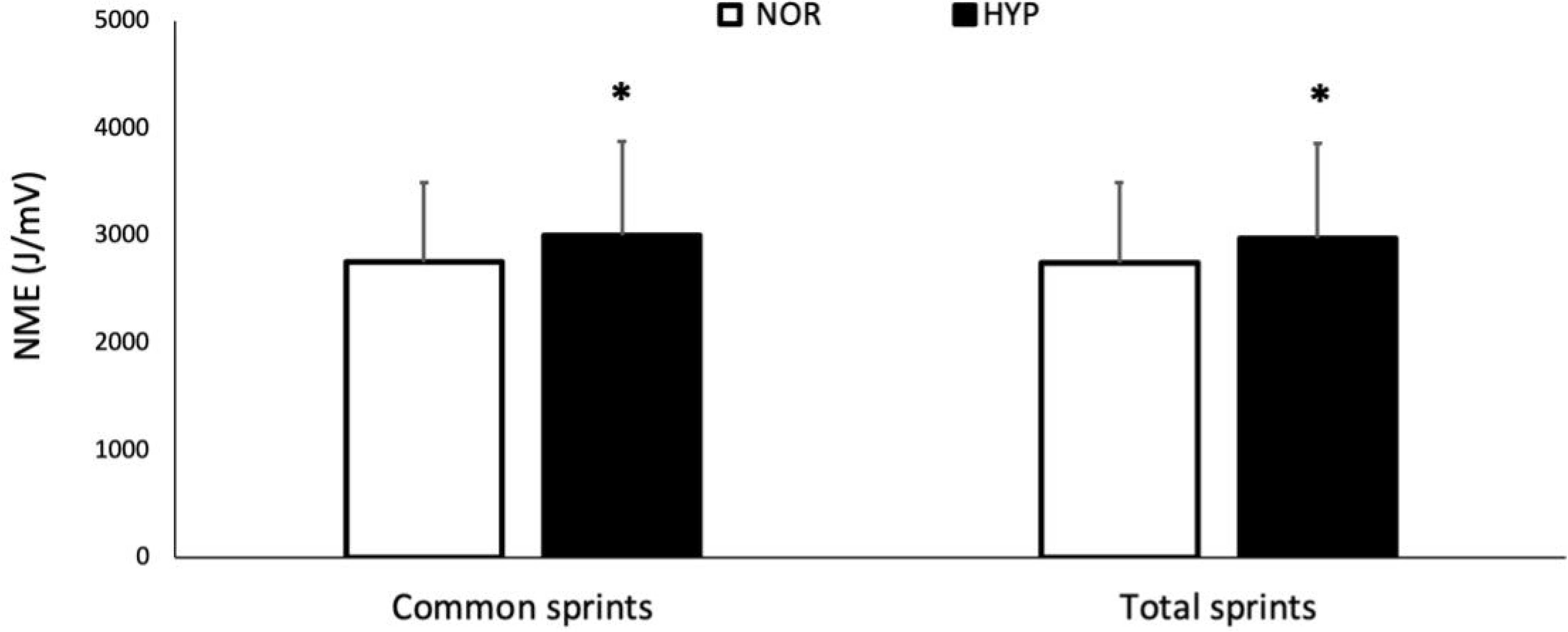
Changes for neuromuscular efficiency (NME) over the entire series of sprints (NME_tot_) and for the same number of sprints performed in both conditions (NME_com_) over the repeated-sprint ability test in normoxia and hyperoxia (F_I_O_2_ 0.40). Data are presented as means ± SD. * Small effect between conditions. NME_tot_ values were 2747.12 ± 746.54 AU in NOR and 2975.51 ± 879.80 AU in HYP. NME_com_ values were 2749.86 ± 746.07 AU in NOR and 3000.14 ± 872.70 AU in HYP.

## 5 Discussion

The present study was performed to assess the acute effects of breathing a hyperoxic mixture on psycho-physiological responses and performance during a repeated-sprint ability test in athletes. The main results were that hyperoxia enhanced sprint endurance during short cycling sprints and the training load of the session. This improvement in mechanical performance may have been due to the concomitant increase in muscle oxygenation fluxes during sprints and recovery periods subsequent to the enhancement in arterial O_2_ saturation and in neuromuscular recruitment.

### 5.1 Hyperoxia and repeated-sprint session training load

The present study demonstrated that the initial sprint mechanical indices (work and PPO) were not altered by HYP. This comes with no surprise since the ability to produce maximal power during short efforts is mainly derived from anaerobic energy sources (Calbet, De Paz, Garatachea, de Vaca, & Chavarren, 2003). This is in keeping with the robust finding that lowering F_I_O_2_ (i.e., hypoxia) also does not impair maximal power output (Balsom, Ekblom, & Sjödin, 1994; Billaut & Buchheit, 2013; Smith & Billaut, 2010). Taken together, this data suggests that short-term maximal performance is well preserved in face of a wide range of F_I_O_2_. The initial sprint performance is related to the mechanical decrement occurring in subsequent sprints (Bishop, Lawrence, & Spencer, 2003), thus we can exclude the influence of this methodological factor on RSA in the present study. When sprints are repeated, the accumulation of metabolic end-products linked to high rates of anaerobic glycolysis induces muscle fatigue. In this regard, breathing more O_2_ may attenuate the loss of power and prolong exercise. Indeed, participants produced ~14% more work over the entire series in HYP compared to room air, and this enhanced performance likely came from a greater aerobic ATP production (Calbet, Martín-Rodríguez, Martin-Rincon, & Morales-Alamo, 2020). Such ergogenic effect has also been reported during sprints of 15 to 30-s duration with 100% O_2_ (Kay et al., 2008; Porter et al., 2019). Our data therefore adds to the literature that mild hyperoxia (40% O_2_) can also enhance RSA during very short efforts.

In addition to the greater performance, HYP exacerbated the breathing and total effort perception. Overall, HYP increased the session TL by ~19%, and this occurred in 12 participants out of 13. A further interesting result, with practical applications for training, was that HYP enhanced performance by increasing the number of repetitions without altering the work performed during the sprints that were common to both conditions. This represents a differing ergogenic impact of hyperoxia during short vs long sprints, in which power output appears to increase during the sprint (Kay et al., 2008; Porter et al., 2020). This is likely caused by the different muscle energetics during efforts of varying duration (Calbet et al., 2020). Since aerobic processes become more predominant as 6-s sprints are repeated (Gaitanos et al., 1993), it is not surprising to observe the ergogenic impact of hyperoxia in the latter stages of a RSA protocol. This difference may also be caused by the mild hyperoxia used in the present study compared with a 1.0 F_I_O_2_ in former studies. Nonetheless, for the coach or sport scientist aiming to increase TL during RSA sessions with team- and racquet-sport athletes, it may be more relevant to use open-loop exercises in which athletes can perform more repetitions before ending a session. Such training approach will have to be tested in long-term, periodizsed, longitudinal studies.

### 5.2 Hyperoxia and arterial and muscle oxygenation

Although hypoxemia is rarely seen during RSA protocols at sea level, it may still develop during longer exercise protocols and hinder performance (Billaut & Smith, 2010). Considering the criteria defined by Dempsey and Wagner (Dempsey & Wagner, 1999), our results show that seven participants experienced mild hypoxemia (>3% S_p_O_2_ fall from resting levels) and one participant experienced moderate hypoxemia (S_p_O_2_ < 93%) during the test. Oxygen availability is generally accepted to play an important role during RSA tests. In fact, arterial hypoxemia imposed by hypoxia induces greater metabolic perturbations and reduces performance during repeated sprints with brief recovery (Balsom, Ekblom, & Sjödin, 1994; Smith & Billaut, 2010; 2012). Here, HYP had a clear positive impact to prevent the fall in S_p_O_2_ and maintained it at 99-100% throughout the test. Interestingly, the five athletes displaying no significant fall in S_p_O_2_ in NOR were those exhibiting the smallest difference in mechanical work between NOR and HYP.

To our best knowledge, the present study is the first to report changes in muscle oxygenation during repeated sprints under hyperoxia. The performance gains in HYP were concomitant with greater muscle oxygenation fluxes. Typically, muscle deoxygenation increases rapidly in the first sprint and then plateaus in the subsequent sprints, which indicates maximal skeletal muscle O_2_ extraction early in the exercise (Billaut & Buchheit, 2013; Buchheit & Ufland, 2011; Buchheit et al., 2009; Smith & Billaut, 2010). HYP could not enhance this mechanism further since both [HHb/Mb]_max_ and TSI_min_ remained similar to those observed in NOR. However, Δ[HHb/Mb]_exercise_ and ΔTSI_exercise_ increased in HYP, indicative of greater peripheral muscle O_2_ extraction when more O_2_ is available before reaching maximal limit. Thus, one may argue that the relative contribution of the oxidative metabolism vs anaerobic glycolysis was greater during the sprints in HYP, thereby contributing to prolong exercise.

This greater aerobic contribution during exercise was probably possible through changes in the recovery. The lower [HHb/Mb]_min_ and greater TSI_max_, Δ[HHb/Mb]_recovery_ and ΔTSI_recovery_ suggest that HYP also enhanced oxygenation status of the muscle during recovery periods. Muscular activity is dependent on the balance between O_2_ delivery and O_2_ consumption by the skeletal muscle fibers, and convective delivery of O_2_ is regulated by blood flow supplying the working muscles and the O_2_ content of the blood. HYP enhanced S_p_O_2_ and also [THb/Mb]_max_ suggesting a greater regional volume of O_2_-rich blood. Although we could not measure blood flow per se, it is still likely that O_2_ delivery and utilisation were enhanced during the recovery between sprints. Muscle reoxygenation plays a critical role on RSA (Billaut & Buchheit, 2013; Buchheit et al., 2009). The present results confirm these findings and together highlight muscle reoxygenation capacity as an important factor of the ability to reproduce mechanical performance in subsequent sprint repetitions. The recovery of phosphocreatine and muscle reoxygenation after exercise presents similar kinetics (Chance, Dait, Zhang, Hamaoka, & Hagerman, 1992; Kime et al., 2003; McCully et al., 1994), and it has been reported that hyperoxia accelerates phosphocreatine resynthesis (Haseler, Richardson, Videen, & Hogan, 1998; Hogan, Richardson, & Haseler, 1999). It is therefore probable that a better muscle reoxygenation in HYP would have translated into a better PCr resynthesis during the recovery periods and a higher PCr availability for subsequent sprints.

Although the maximal peripheral O_2_ uptake during the sprints did not appear to differ between the environments, probably due to the very high intensity of the efforts and stress put on the oxidative pathway, HYP still generated marked amplitudes of tissue oxygenation during both sprint and recovery. Repeated fluctuations of tissue partial pressure of O_2_ are necessary for muscular oxidative capacity adaptations (Casey & Joyner, 2012; Daussin et al., 2008). Thus, in addition to increasing a session TL, hyperoxia may also be an effective environmental stress for inducing peripheral adaptations by imposing repeated muscle oxygenation fluxes.

### 5.3 Hyperoxia and muscle recruitment

Under substantial fatigue (evaluated at >10% fatigue index or sprint decrement), a concomitant attenuation of EMG activity and mechanical performance is usually observed across sprint repetitions (Billaut & Buchheit, 2013; Mendez-Villanueva, Hamer, & Bishop, 2008; Mendez-Villanueva, 2007; Racinais et al., 2007). This suggests that the ability to fully activate the working muscles may be incriminated in the loss of mechanical output. Motor neuron activity is greatly influenced by O_2_ availability (Amann, Romer, Subudhi, Pegelow, & Dempsey, 2007; Bigland-Ritchie, Dawson, Johansson, & Lippold, 1986; Szubski, Burtscher, & Löscher, 2006). In the RSA literature, the reduction in S_p_O_2_ has been correlated with the attenuation in surface EMG activity (Billaut & Smith, 2010), and has also been shown to accentuate the decline in central motor command assessed from peripheral muscle stimulation (Billaut et al., 2013). Our results therefore suggest that O_2_ supplementation during sprints helps maintain a higher muscle recruitment (as evidenced by a ~10% higher sum-iEMG indices in HYP). The most likely explanation would be that the greater tissue perfusion and oxygenation improved the metabolic milieu of contracting muscles and attenuated the reflex inhibition originating from group III and IV afferents, thereby maintaining neural drive to skeletal muscles (Amann & Dempsey, 2008).

There is also the possibility that changes in EMG indices related at least in part to sarcolemma excitability and changes in conduction velocity of action potentials. Membrane excitability is impaired during intense fatiguing exercise as a result of a lower activity of the sodium(Na^+^)/potassium(K^+^)-adenosine triphosphatase (NKA) activity to maintain transmembrane ionic gradient caused by the decline in pH and accumulation of inorganic phosphates. Greater muscle perfusion and oxygenation during sprints could have altered energy metabolism and the subsequent accumulation of by-products and acidosis that resulted in lesser sarcolemma dysfunction (Linossier et al., 2000). Future studies should more robustly investigate the potential attenuation of centrally mediated decline in neural drive and alterations in M-wave amplitude.

## 6 Conclusion

This study demonstrated that breathing a hyperoxic mixture with 40% O_2_ improved the total work performed and the associated training load during an open-loop test of RSA in trained team- and racquet-sport athletes. This ergogenic effect occurred concurrent with an increase in muscle oxygenation fluxes during sprints and recovery periods and in neuromuscular recruitment. Although this study showed promising results about the effectiveness of mild hyperoxia to enhance TL during RSA sessions, important factors such as manipulation of F_I_O_2_ and optimum exercise-to-rest ratios remain to be explored for efficient applications.

## 7 Author Contributions

SCK and FB conceptualized and designed the research project; SCK acquired the data and conducted the statistical analysis; SCK interpreted results with assistance from FB; SCK wrote the manuscript with revisions from FB; all authors reviewed and agreed upon the final manuscript.

## 8 Acknowledgments

The authors thank the athletes for their participation in this study. We also sincerely thank all graduate and undergraduate students from our research group for their valuable help, as well as TMC distribution for the generous loan of the oxygen concentrator.

